# Enhanced Activity of Apramycin and Apramycin-Based Combinations Against *Mycobacteroides abscessus*

**DOI:** 10.1101/2025.04.09.648020

**Authors:** Yanqin Huang, Katherine A. Truelson, Isabella A. Stewart, George A. O’Doherty, James E. Kirby

**Affiliations:** Department of Pathology, Beth Israel Deaconess Medical Center, Boston, MA, USA; Harvard Medical School, Boston, MA, USA; Department of Chemistry, Northeastern University, Boston, MA 02115, USA

**Keywords:** *Mycobacterium abscessus*, synergy, checkerboard, apramycin, time-kill, amikacin, clofazimine

## Abstract

**Background:** *Mycobacteroides abscessus* are rapidly growing non-tuberculous mycobacteria that cause chronic lung and soft tissue infections. Treatment options are often severely limited due to intrinsic resistance to most antimicrobials. Amikacin has historically been a mainstay of combination treatment regimens. However, irreversible hearing loss and vestibular toxicity have led to a search for alternative agents. Apramycin is a novel aminoglycoside currently in phase I clinical trials that may offer lower potential for ototoxic and renal toxic side effects.

**Objectives:** The goal of this study was to compare apramycin’s *in vitro* activity with amikacin and other aminoglycosides against a large collection of *M. abscessus* clinical isolates, both alone and in combination with clofazimine or linezolid. We also tested the activity of apramycin against a more limited collection of other species of rapidly growing mycobacteria.

**Methods:** Analysis was performed using reference broth microdilution minimal inhibitory concentration testing, inkjet printer-assisted checkerboard assays, and time-kill assays.

**Results:** Against *M. abscessus*, the MIC_50/90_ for apramycin (2 µg/mL) was 8-fold lower than for amikacin (16 µg/mL). Plazomicin was inactive, and organisms were rarely susceptible to tobramycin. Synergy was not detected by checkerboard assay. In time-kill studies, clofazimine modestly potentiated activity of apramycin and. to a lesser extent, amikacin. Apramycin and amikacin showed delayed bacterial killing that either achieved or approached a bactericidal threshold. Apramycin was similarly potent against other rapidly growing mycobacteria tested.

**Conclusions:** Apramycin exhibits more potent *in vitro* activity against a diverse set of *M. abscessus* and other rapidly growing mycobacteria than approved aminoglycosides.

## Introduction

*Mycobacteroides abscessus* are rapidly growing mycobacterium that cause chronic progressive lung and soft tissue infections.^1^ They are often multidrug-resistant; prolonged treatment courses are required for suppression, control, or cure when possible, leading to cumulative toxicities. The cure rate for lung infection is ∼30–50% with surgical intervention often needed for infection site control.

Currently combinatorial therapy includes choice of two to five antimicrobials guided by antimicrobial susceptibility testing. Amikacin is one of the recommended parenteral agents for initial combination therapy in updated ATS/ERS/ESCMID/IDSA Clinical Practice Guideline guidelines for *M. abscessus* lung infection.^2, 3^ However, there is no consensus on length of amikacin treatment; in practice, parental agents such as amikacin are often used for a month or longer.^2^

Unfortunately, the toxicities of extended aminoglycoside treatment include irreversible vestibular and cochlear damage, and renal toxicity. Apramycin is a novel aminoglycoside with a mono-substituted 2-deoxystreptamine attached to a bicycle sugar containing disaccharide having a shifted binding site in the 30S ribosome compared with amikacin. In a human phase I clinical trial (NCT04105205) apramycin was safe (results not published); and in the rat model, renal toxicity appeared to be much lower than for gentamicin.^4^ In an explanted cochlear model, apramycin did not appear to be associated with ototoxicity, potentially because of lower affinity for mitochondrial and/or eukaryotic cytoplasmic ribosomes. ^4-10^ It is now being studied in a second phase I trial in the US (NCT05590728) to determine lung epithelial lining fluid and serum pharmacokinetics.

An early disk diffusion study found that apramycin was active *in vitro* against *M. abscessus* clinical isolates; however, quantitative minimal inhibitor concentration data were not assessed.^11^ A more recent study found apramycin minimal inhibitory concentration (MIC) values of 0.5 µg/mL for six *M. abscessus* clinical isolates — two isolates each of subsp. *abscessus, bolleti*, and *massiliense*.^12^ In this limited analysis, apramycin was found to have lower MICs than amikacin. It also was found in contrast to amikacin to be bactericidal in time-kill study analysis for four *M. abscessus* isolates examined. Genetic experiments support the role of the multi-acetyltransferase, Eis2, found ubiquitously in *M. abscessus* as the direct cause of curtailed amikacin activity relative to apramycin, as only amikacin is a substrate for this resistance enzyme.^12-14^ In addition, apramycin activity unlike that of amikacin was not affected by the transcriptional regulator, WhiB7, which upregulates a set of genes conferring resistance to several frontline antimicrobial therapies, in the case of amikacin through upregulation of Eis2.^15^

*In vivo* experimental evidence suggests that apramycin is a promising treatment for lung and soft tissue infections, the major sites of *M. abscessus* disease.^12, 16^ Murine studies showed that apramycin lung epithelial lining fluid levels approximate those in plasma. In turn, apramycin proved efficacious in treatment in murine pneumonia models of infection caused by *Klebsiella pneumoniae, Acinetobacter baumannii, Pseudomonas aeruginosa*, and *M. abscessus*. Apramycin was also shown active in thigh and urinary tract models.^4, 17^ Modeling indicates a high likelihood of apramycin target attainment in humans dosed at 30mg/kg dose and a pathogen MIC ≤ 8 µg/mL in pneumonia and urinary tract gram-negative ESKAPE pathogen infection,^16^ providing a potential categorical breakpoint for *M. abscessus* susceptibility.

Taken together, existing data suggest that apramycin might have more potent activity and lower toxicity compared with amikacin for *M. abscessus* infection. This current work significantly extends activity spectrum analysis of apramycin in comparison with other aminoglycosides against contemporary clinical isolates *M. abscessus* and other rapidly growing mycobacterium species, and tests apramycin’s potential *in vitro* synergistic activity in combination with other antimicrobials used for *M. abscessus* treatment.

## Methods and Materials

### Antibiotics

Apramycin sulfate (Alfa Aesar, MA, USA, Cat# J66616, lot#Y05F120); clofazimine (Acros Organics, Geel, Belgium, Cat# 461760010, lot# A0395020); linezolid (Acros Organics, Geel, Belgium, cat# 460592500, lot# A0400349); tobramycin sulfate (Research Products International, IL, USA, cat#T45000-1.0, lot#10298-155238); plazomicin sulfate (ToKu-E, WA, USA cat #P140, lot#P140-01US); and amikacin disulfate (Alfa Aesar, cat#J63862.14, lot# N08H025) were obtained from indicates sources

### Bacterial isolates

Clinical isolates were from Beth Israel Deaconess Medical Center (Boston, MA, USA) from years 2016-2023 under an Institutional Review Board-approved protocol and from the American Type Culture Collection (Manassas, VA) as listed in **Table S1**. All isolates and strains were stored at -80°C and minimally passaged until use. Subspeciation was determined by a previously described multi-plex PCR typing method^18^ and/or by the University of Tyler, TX, Mycobacteriology and Nocardia Reference Laboratory.

### Minimal inhibitory concentration testing

Broth microdilution antimicrobial susceptibility testing was performed as recommended by the Clinical Laboratory and Standards Institute (CLSI)^19, 20^. Serial two-fold dilutions of freshly prepared drugs were dispensed using the HP D300 digital dispensing system (HP, Inc., Palo Alto, CA) into sterile, round-bottom, polystyrene 96-well plates (Evergreen Scientific, Los Angeles, CA) to achieve final desired antimicrobial concentrations as we previously described.^21-25^ *M. peregrinum* ATCC 700686 and *Escherichia*.*coli* ATCC 25922 were used as quality control strains.^26^ Following incubation at 30°C in ambient air, minimal inhibitory concentration (MIC) results were recorded from day 3 to day 5 per CLSI recommendations.

### Checkerboard Synergy Assays

Antimicrobials combinations of apramycin or amikacin with linezolid and clofazimine were dispensed using the HP D300 digital dispensing system (HP, Inc., Palo Alto, CA) as previously described in full checkerboard synergy arrays ^21, 27^ and tested against three *M. abscessus* clinical isolates, NTM18, NTM27, and NTM28, in duplicate. The fractional inhibitory concentration index (FICI) was determined by summing the individual FICs for each antibiotic in each inhibited well where the FIC for each antimicrobial equals the inhibitory concentration of each antimicrobial in the combination divided by the MIC of the antimicrobial when tested alone. The FICI is the lowest summation for which complete visual growth inhibition was observed. The highest (most conservative) FICI among replicates was scored as indicating synergy (≤ 0.5), indifference (0.5 < FICI ≤ 4.0), or antagonism (FICI > 4.0), respectively.^28^

### Time-Kill Assays

Time-kills assays were performed in duplicate, as previously described, against NTM27 and NTM28 with a starting inoculum of ∼10^6^ CFU/mL.^2^ Apramycin or amikacin was tested either alone or combination at indicated multiples of MICs determined by broth microdilution. Bacteria were incubated at 30ºC with sampling for CFU determination at indicated time points. Bactericidal activity was defined as a 3log_10_ reduction in viable counts compared with the initial inoculum after the incubation period indicated. Synergy was defined as a 2log_ 10_ drop in an antimicrobial combination at 72 h compared with the effects of either antibiotic tested alone at the same concentrations.

### Spontaneous resistance frequency

Log-phase NTM27 and NTM28 *M. abscessus* susp. *abscessus* were plated at 10^9^, 10^10^ or 10^11^ CFU onto cation-adjusted Mueller-Hinton agar plates containing apramycin or amikacin at a concentration four-fold or eight-fold greater than the antibiotic specific MIC. The resistance frequency was calculated by determining the fraction of bacteria growing on antibiotic versus non-antibiotic containing medium after 5 days of incubation at 30C°.^29^

## Results

### Minimal inhibitory concentration (MIC) data

The MIC_50,_ MIC_90_, and MIC ranges for antimicrobials tested are listed in **Table 1**. The *M. abscessus* isolate subspecies, specimen source, and individual MIC values for each isolate are listed in **Table S1**. Notably, the MIC_50_ and MIC_90_ for apramycin was 8-fold lower than for amikacin. Plazomicin was inactive at the highest concentration tested, while only 12.8% of the isolates were predicted to be susceptible to tobramycin. A smaller number of *M. fortuitum* and *M. chelonae* were also tested with similar apramycin activity and improved amikacin activity compared with *M. abscessus* (**Table S2**).

**Table 1.**
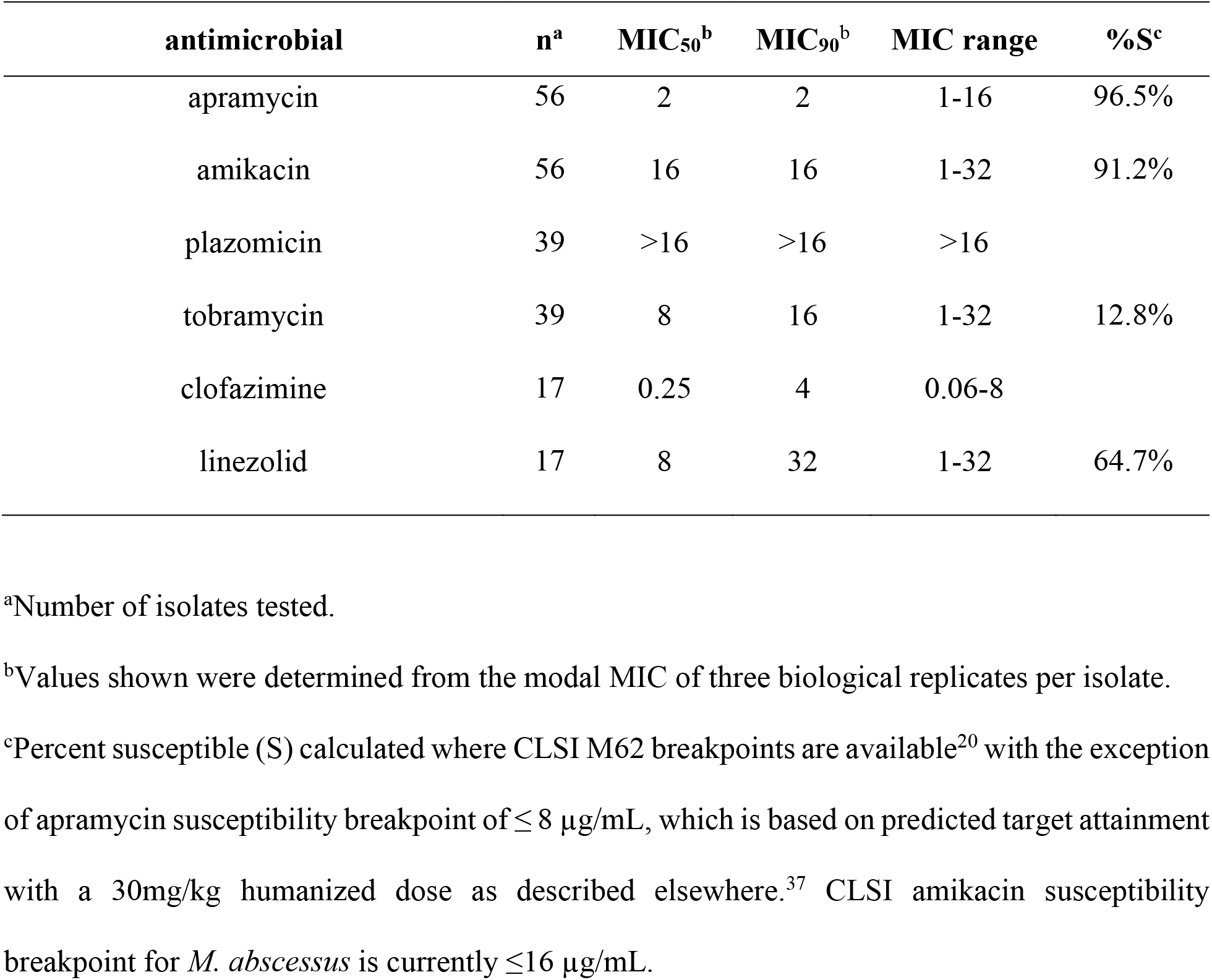
MIC_50_, MIC_90_, and MIC range in µg/mL for aminoglycosides and antimicrobials used in synergy testing.

| antimicrobial | n<sup>a</sup> | MIC<sub>50</sub><sup>b</sup> | MIC<sub>90</sub><sup>b</sup> | MIC range | %S<sup>c</sup> |
| --- | --- | --- | --- | --- | --- |
| apramycin | 56 | 2 | 2 | 1-16 | 96.5% |
| amikacin | 56 | 16 | 16 | 1-32 | 91.2% |
| plazomicin | 39 | >16 | >16 | >16 |  |
| tobramycin | 39 | 8 | 16 | 1-32 | 12.8% |
| clofazimine | 17 | 0.25 | 4 | 0.06-8 |  |
| linezolid | 17 | 8 | 32 | 1-32 | 64.7% |

<sup>a</sup>Number of isolates tested.
<sup>b</sup>Values shown were determined from the modal MIC of three biological replicates per isolate.
<sup>c</sup>Percent susceptible (S) calculated where CLSI M62 breakpoints are available<sup>20</sup> with the exception of apramycin susceptibility breakpoint of ≤ 8 µg/mL, which is based on predicted target attainment with a 30mg/kg humanized dose as described elsewhere.<sup>37</sup> CLSI amikacin susceptibility breakpoint for *M. abscessus* is currently ≤16 µg/mL.

### Checkerboard Synergy Testing

FICI values for combinations of apramycin or amikacin with clofazimine and linezolid against three representative clinical isolates are shown in **Table 2**. Indifference was observed for all combinations against all isolates.

**Table 2.** FICI values for checkerboard testing of apramycin or amikacin in combination with clofazimine or linezolid.

|  | Clofazimine | Linezolid |
| --- | --- | --- |
| Apramycin | 2, 1.5, 1.75<sup>a</sup> | 1.5, 1.5, 1.5 |
| Amikacin | 2, 3, 2.5 | 1.5, 3, 2 |

<sup>a</sup>FICI values, separated by commas, are listed separately for *M. abscessus* subsp. *abscessus* clinical isolates NTM18, NTM27, and NTM28, respectively.

### Time-kill studies

Time**-**kill analysis for comparison of effects of apramycin and amikacin alone and in combination with clofazimine was performed against two representative *M. abscessus* subsp. *abscessus* clinical isolates **(Fig. 1)**. Apramycin, but not amikacin, at 2× MIC reduced CFU at 72 hours by >3 log_10_ for NTM27, but not for NTM28, compared with untreated controls. Clofazimine alone was not fully bacteriostatic or without effect at 2x its MIC. However, at this concentration, it appeared to modestly potentiate the activity of apramycin, as a ≥3 log_10_reduction in CFU compared to untreated controls was now also observed at 1×MIC for NTM27 and at 4× MIC for NTM28. However, neither aminoglycoside was synergistic with clofazimine at 2×MIC, based on the standard threshold criterion of a ≥2 log_10_ decrease in CFU/mL in combination testing compared with the most active single agent. In experiments with extended 5-day incubations, both apramycin and amikacin at 1× and 2×MIC demonstrated continued, slow killing of NTM27, with apramycin achieving bactericidal activity and amikacin approaching this threshold (**Fig. 2**). The combination of either apramycin or amikacin at 2×MIC with clofazimine at 2×MIC was bactericidal, achieving a ≥4 log_10_kill by day 5, effectively sterilizing cultures to the limit of detection for NTM27. NTM28 was not tested in extended time-kill experiments.

**Figure 1.**
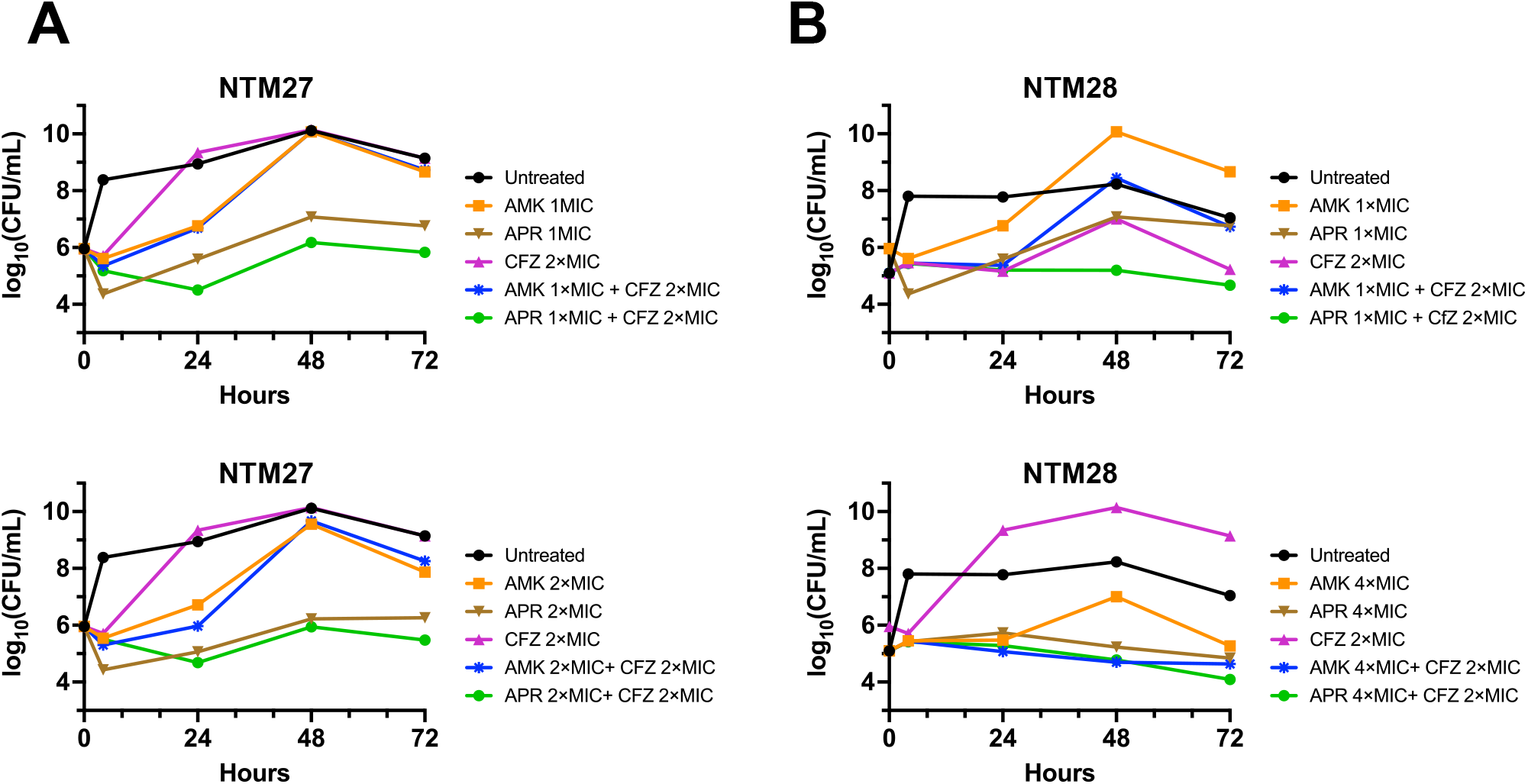
Time-kill analysis of amikacin and apramycin alone and in combination with clofazimine against two *M. abscessus* subsp. *abscessus* clinical isolates. **(A)** Time-kill analysis against strain NTM27 using concentrations of antimicrobials at indicated multiples of MIC values determined by the reference broth microdilution method. **(B)** Time-kill analysis for NTM28. For NTM27, MICs for AMK, APR, and CFZ were 8, 2, and 0.25 µg/mL, respectively. For NTM28, MICs for AMK, APR, and CFZ were 8, 2, and 04 µg/mL, respectively. Abbreviations: AMK, amikacin; APR, apramycin; CFZ, clofazimine.

**Figure 2.**
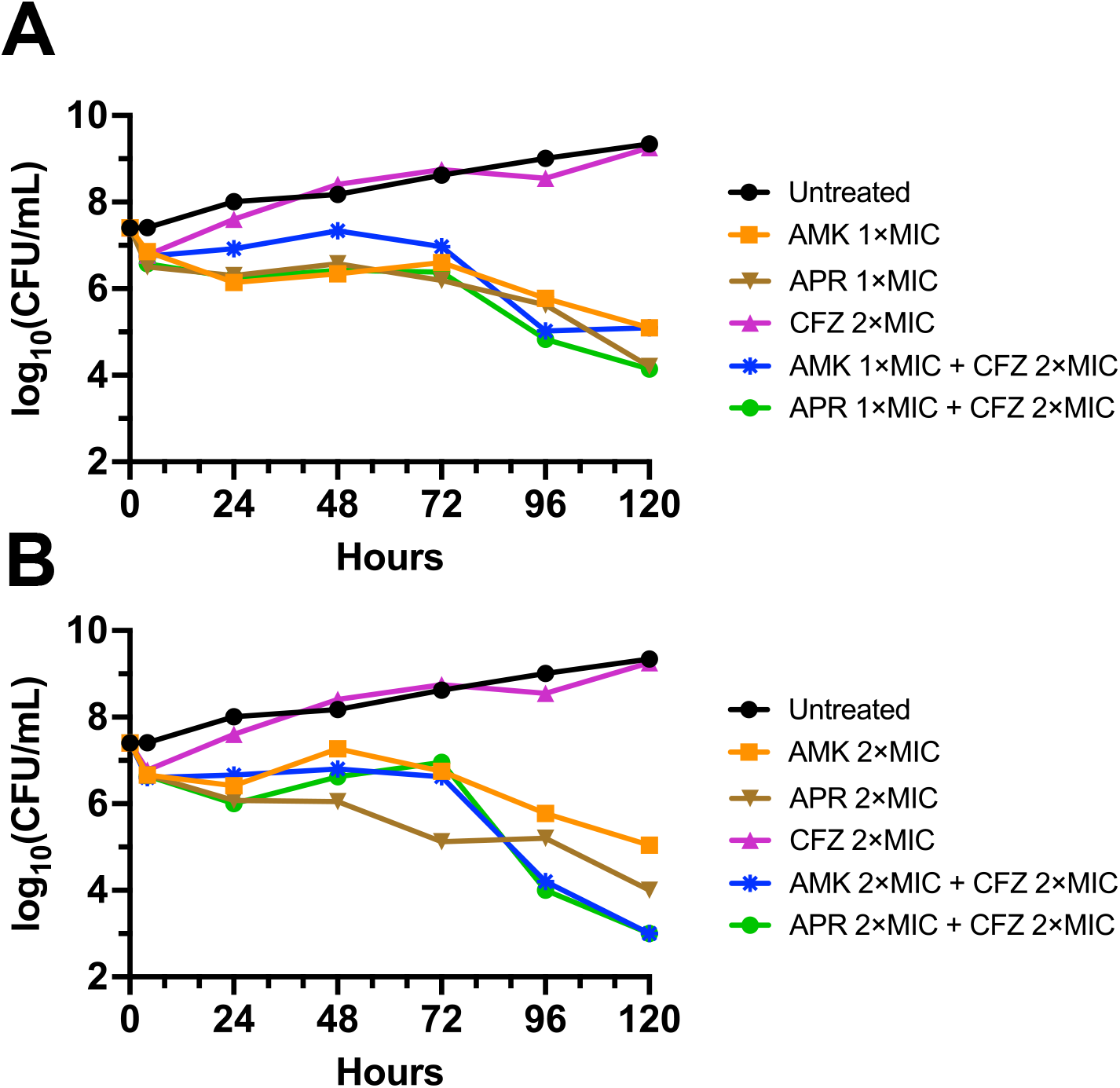
Extended incubation reveals slow bactericidal killing by apramycin and amikacin, modestly potentiated by clofazimine. Time-kill analysis during extended incubation was performed for NTM27 *M. abscessus* susp. *abscessus* with antibiotic concentrations at indicated multiples of MIC values determined by reference broth microdilution. **(A)** Amikacin and apramycin added at 1×MIC alone or in combination with clofazimine. **(B)** Amikacin and apramycin added at 2×MIC alone or in combination with clofazimine. Abbreviations: AMK, amikacin; APR, apramycin; CFZ, clofazimine.

### Spontaneous resistance mutation frequency

Both apramycin and amikacin at 4×MIC and 8×MIC showed a low and essentially identical spontaneous resistance frequency against two *M. abscessus* susp. *abscessus* clinical isolates (**Table 3**).

**Table 3.** Spontaneous resistance frequency in representative *M. abscessus* susp. *Abscessus* isolates at indicate multiples of amikacin and apramycin MIC.

| Strain | Amikacin (MIC, 8µg/mL) |  | Apramycin (MIC, 2 µg/mL) |  |
| --- | --- | --- | --- | --- |
|  | 4×MIC | 8×MIC | 4×MIC | 8×MIC |
| **NTM27** | 4×10<sup>-9</sup> | 2×10<sup>-9</sup> | 2×10<sup>-9</sup> | 2×10<sup>-9</sup> |
| **NTM28** | 4×10<sup>-9</sup> | 7×10<sup>-10</sup> | 4×10<sup>-9</sup> | 3×10<sup>-10</sup> |

## Discussion

Amikacin has long been an integral part of treatment regimens for *M. abscessus* infection. This general bactericidal property of aminoglycosides is thought to provide advantage in rapid clearance of organisms, especially at sites where efficacy of cell-mediated immunity may be limited.

However, there is concern that current breakpoints for amikacin may be set too high for lung infections. Notably, the amikacin susceptibility breakpoint for *Enterobacterales* was recently lowered by CLSI to ≤4 µg/mL, reflecting a more current understanding of pharmacokinetic/pharmacodynamic (PK/PD) relationships^30^ significantly lower than the 16 µg/mL susceptibility breakpoint for *M. abscessus*.^12, 31, 32^ Furthermore, results from hollow fiber infection models suggest that dosing regimens required for reliable efficacy would invariably be associated with ototoxicity.^31^ In the present study, we found that the MIC_50_/MIC_90_ for amikacin across a large set of clinical isolates was 16 µg/mL, suggesting that amikacin should offer limited clinical benefit for the direct treatment of pulmonary airway disease. Yet, there is still evidence of clinical efficacy in uncontrolled retrospective studies comparing different multidrug regimens with or without amikacin, despite PK/PD parameters that would typically suggest limited utility.^33, 34^ Overall, however, amikacin appears far from an ideal choice in regimens where bactericidal activity is desired.

Therefore, a more potent, bactericidal antibiotic without the limiting side effects of amikacin would be welcome. Here, we found that apramycin was 8-fold more active (MIC_50_/MIC_90_) than amikacin by weight. Recently, in a murine lung infection model, a human-equivalent apramycin dose of 30 mg/kg was found to offer a 99% probability of achieving a 2-log_10_reduction of *A. baumannii* for isolates with an MIC ≤16 µg/mL.^35^ This MIC threshold, if found to be similarly applicable to *M. abscessus* infection, would encompass 100% of the largest reported set of *M. abscessus* clinical isolates examined to date.

In contrast to apramycin, tobramycin was generally inactive, with MICs roughly correlating with amikacin MIC values, while plazomicin was universally inactive. The *M. abscessus* genome encodes an AAC(2′) enzyme sharing 35% identity and 53% similarity with the aminoglycoside *N*-2′-acetyltransferase-Ia [AAC(2′)-Ia] from *Providencia stuartii*, which is known to inactivate plazomicin and tobramycin through acetylation of the *N*-2′ position of their 4-*O*-linked sugars.^14^ The alpha-fold-predicted structure of the *M. abscessus* protein is highly similar to the *Providencia* enzyme (PDB 6VRO) with a compelling root mean squared deviation of 0.8 Å (**Fig. S1**). Therefore, this enzyme is a prime candidate for inactivation of plazomicin in *M. abscessus*. However, we have not confirmed this hypothesis through genetic knockout experiments as this was not a specific focus of the current work.

The lack of formal synergy between apramycin or amikacin and clofazimine or linezolid examined in checkerboard testing (also observed in time-kill studies for the combination of aminoglycosides and clofazimine) suggests that subtherapeutic concentrations of either aminoglycoside (below those suggested by pharmacokinetic/pharmacodynamic relationships to be efficacious as single agents) should not be relied upon to provide benefit in combination treatment, which is a significant drawback for amikacin due to its much higher minimum inhibitory concentration levels. However, the results also suggest some enhanced killing with clofazimine-apramycin combinations, which may translate to improved therapeutic efficacy. A similar rate of spontaneous resistance to apramycin and amikacin, consistent with an overlapping, 16S rRNA target in this species (**Fig. S2**), suggests that resistance to apramycin is unlikely to develop more rapidly than amikacin during treatment. However, for both drugs, the presence of a single-copy *M. abscessus* rRNA operon highlights the additional necessity of combination treatment to prevent spontaneous, single-step resistance leading to treatment failure.^36^

In conclusion, apramycin shows significant promise as a potent, bactericidal agent against the vast majority of *M. abscessus* isolates. Our findings support further investigation of apramycin for the treatment of *M. abscessus* infections either by intravenous or nebulized administration routes.

## Funding

This work was supported by a Novel Therapeutics Delivery Grant from Massachusetts Life Science Center to J.E.K. Y.H. was supported in part by a National Institute of Allergy and Infectious Diseases training grant (T32AI007061) and an Academy of Clinical Laboratory Physicians and Scientists (ACLPS) Paul E. Strandjord Young Investigator Research Grant. The content is solely the responsibility of the authors and does not necessarily represent the official views of the NIH. The HP D300 digital dispenser used in these studies was provided by TECAN (Morrisville, NC). TECAN had no role in study design, data collection, or interpretation.

### Transparency Declaration

None to declare.

