## Supplementary Figures for "Enhanced Activity of Apramycin and Apramycin-Based Combinations Against *Mycobacteroides abscessus*"

a

[a]Department of Pathology, Beth Israel Deaconess Medical Center, Boston, MA, USA

[b]Harvard Medical School, Boston, MA, USA

[c]Department of Chemistry, Northeastern University, Boston, MA 02115, USA

Running Head: Apramycin combinations against *M. abscessus*

### Supplementary Figures

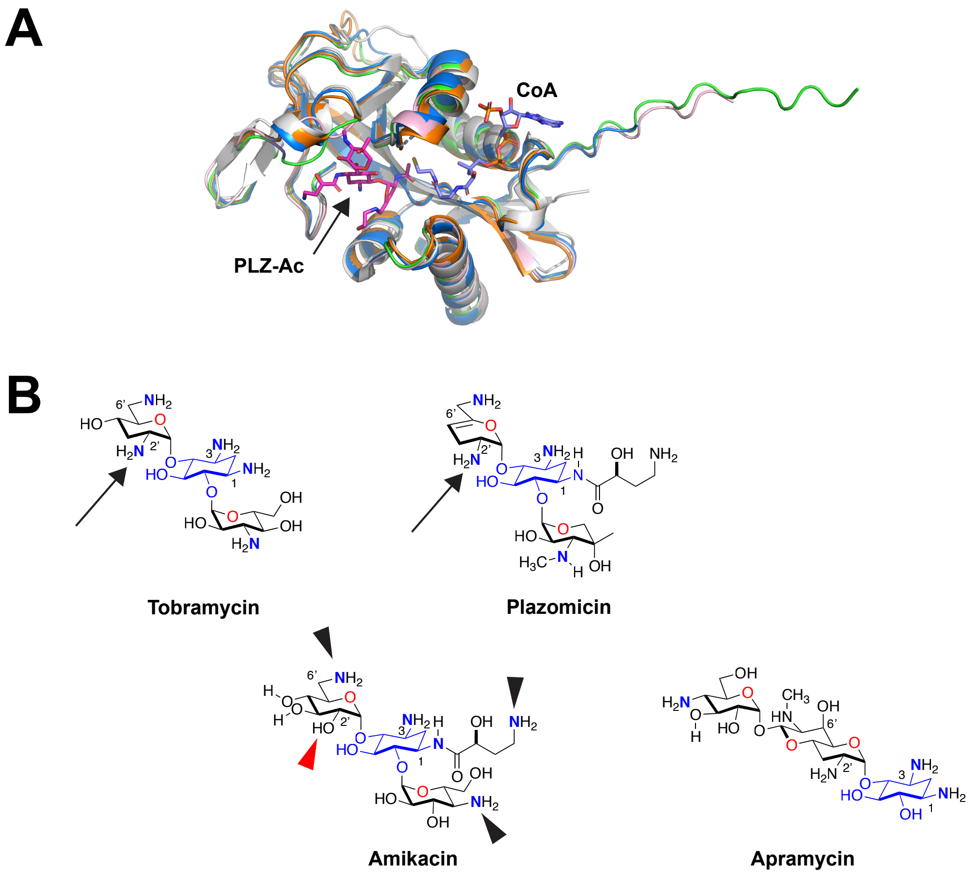


**Figure S1. Inactivation of aminoglycosides in *M. abscessus*. (A)** The structures of *M. abscessus* orf 4395 (a putative AAC(2') acetylase) and homologs from *M. fortuitum*, *M. chelonae*, and *M. tuberculosis* were predicted by ColabFold v1.5.5: AlphaFold2 using MMseqs2 [1]. The highest ranked structures from each alpha-fold prediction were aligned with the *P. stuartii* aminoglycoside *N*-2'-acetyltransferase-Ia (AAC(2')-Ia) structure, PDB 6VOU [2]. Shown are an overlay of aligned AAC(2')-Ia (gray) from the *P. stuartii* structure with acetylated plazomicin (PLZ-Ac, magenta) and coenzyme A (CoA, blue) with predicted protein homologues from *M. abscessus* ATCC 19977 (Mab Orf 4395, green); *M. fortuitum* DSM 46621 (orange); *M. chelonae* ATCC 35752 (light pink); and *M. tuberculosis* H37Rv (blue). RMSD for all aligned pairwise comparisons with the *P. stuartii* enzyme were < 0.9 Å. Alignments and graphical representations were performed using Pymol version 3.1.3. **(B)** Chemical structures of apramycin, amikacin, plazomicin, and tobramycin rendered in ChemDraw version 22.2.0.3348. Positions in tobramycin and plazomicin presumptively modified by the AAC(2')-Ia homologue are indicated by arrows. Amikacin has a hydroxyl group in place of the primary amine at the *C*-2′ position (red dart), which eliminates this antibiotic as a substrate for AAC(2′)-Ia-mediated modification. The indicated primary amines in amikacin (black arrowheads) are identified sites of *N*-acetylation by the Eis enzyme from *M. tuberculosis* [3], which is highly homologous to the Eis2 enzyme in *M. abscessus* (alignment RMSD = 1.4 Å). Eis2 has been found responsible for the elevated amikacin MICs in the latter pathogen. [4, 5]

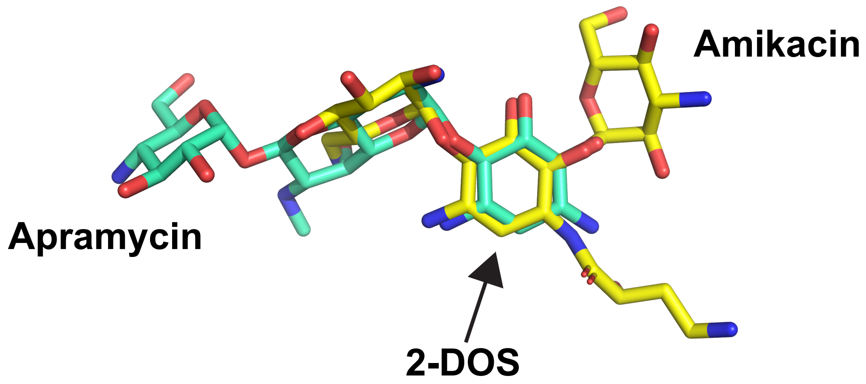


**Fig S2. Overlap of apramycin and amikacin binding sites in the prokaryotic ribosome.** Alignment and overlay of PDB 7PJS (apramycin, green) and 8SYL (amikacin, yellow). Both antibiotics bind to helix 44 of the 16S rRNA in the 30S ribosomal subunit. Their respective 2-deoxystreptamine (2-DOS) rings superimpose (arrow). Amikacin interacts with 16S rRNA G1405. In contrast, apramycin is shifted away from G1405, allowing it to make unique contacts with prokaryotic helix 44 in a region divergent from the corresponding mitochondrial ribosome structure, which may account for apramycin's more favorable side effect profile.
